# Highly Efficient Knockin in Human iPS Cells and Rat Embryos by CRISPR/Cas9 Molecular Optimization

**DOI:** 10.1101/2021.09.03.458921

**Authors:** Vanessa Chenouard, Isabelle Leray, Laurent Tesson, Séverine Remy, Agnès Fortun, Karine Bernardeau, Yacine Cherifi, Laurent David, Ignacio Anegon

## Abstract

The CRISPR/Cas9 system is now the gold standard for the generation of genetically modified cell and animal models but knockin is a bottleneck. One reason could be that there is no consensus regarding the concentrations of its components to be used. Here, we defined optimal Cas9 protein, guide RNA and short donor DNA concentrations on a GFP to BFP conversion model of human induced pluripotent stem cells and point mutations on rat transgenic embryos. With a molecular rational approach of the CRISPR/Cas9 system and study of ribonucleoprotein complex formation by nanodifferential scanning fluorimetry, we defined that Cas9/guide RNA 1/1 molar ratio with 0.2μM and 0.4μM of Cas9, coupled with 2μM of ssODN are sufficient for optimal and high knockin frequencies in rat embryos and human induced pluripotent stem cells, respectively. These optimal conditions use lower concentrations of CRISPR reagents to form the RNP complex than most conditions published while achieving 50% of knockin. This study allowed us to reduce costs and toxicity while improving editing and knockin efficacy on two particularly key models to mimic human diseases.

## I. Introduction

Genetically modified models are necessary tools to better understand gene functions and study human diseases. Both animal and cellular models offer pros, cons and hurdles that make them complementary. In particular, for genetically modified animal models, the rat has always been a model of choice to study gene functions or model human diseases but suffered from a lack of accessible tools for decades (Chenouard et al. 2021). In parallel, cell models have been transformed fifteen years ago by the development of human induced pluripotent stem (hiPS) cells (Takahashi and Yamanaka 2006; Yu et al. 2007), which offer the flexibility necessary to mimic most human diseases. Both models have been revolutionized by the conversion of CRISPR/Cas bacterial immune systems into genome editing toolbox (Jinek et al. 2012) to precisely create almost any knockout (KO) or knockin (KI) models (single or multiplex KO, point mutation, conditional KO, reporter…). Nevertheless, efficacy, at least in the most complex KI cases with multiple or large DNA donors DNA, is still too low for an efficient development.

The CRISPR/Cas system is composed of a Cas9 nuclease able to specifically cleave a target DNA sequence by the guidance of an RNA molecule after formation of a ribonucleoprotein (RNP) complex. The concentrations of Cas9 and guide RNA (gRNA) for RNP formation used to deliver into hiPS cells or rodent embryos are quite heterogenous in the literature (Skarnes et al. 2019; Xu et al. 2019; Kaneko and Nakagawa 2020; Remy et al. 2017). For instance, RNP molar ratio (Cas9/gRNA) of 1/5 (Skarnes et al. 2019) have been electroporated to hiPS cells while others used 1/1,3 ratio (Xu et al. 2019). For rat embryos, 1/26 molar ratio (Kaneko and Nakagawa 2020) has been used whereas us and others have used 1/1.6 ratio (Remy et al. 2017). Moreover, conditions were described with mass or massic concentration and required calculations to compare between experiments. The mass of one molecule of DNA depends on its size which complicates comparison of massic conditions except for similar donors. To circumvent this, it would be more appropriate to work in molarity rather than mass to facilitate comparison between studies and repetition of experiments in other laboratories.

In our study, we analyzed the basic molecular aspects of this system to define Cas9 and dual gRNA (dgRNA) lowest but most efficient concentration for optimal RNP (Cas9/dgRNA) formation and the optimal single-stranded oligonucleotide (ssODN) concentration for the electroporation of hiPS cells and rat embryos to generate KI models. We first challenged the system with electroporation of different concentrations of Cas9 and Cas9/dgRNA/ssODN ratios into hiPS cells. They permanently express GFP and KI converted them into BFP to facilitate the readout by flow cytometry. We then checked RNP complex formation efficiency *in vitro* by nano differential scanning fluorimetry (NanoDSF) to define the best RNP (Cas9/dgRNA) ratio. This ratio was finally applied *in vivo* on the hiPS cells GFP to BFP conversion model and rat embryos point mutations model. It improved the generation of KI in these two particularly key models of human diseases. In turn, those optimized conditions reduced the operating costs of such experiments.

## II. Results

### Molecular optimization of GFP to BFP conversion on hiPS cells

Biallelic homozygous GFP iPS cells were converted into BFP expressing cells by CRISPR/Cas9 KI using a BFP ssODN **(Fig. 1A)** and phenotyped by flow cytometry to define different cell populations that lost GFP expression and/or acquired BFP expression as a consequence of NHEJ or KI, respectively **(Fig. 1B)**. We could define cell subsets that still expressed GFP (presumably with both or one GFP allele conserved), that had completely lost GFP expression (presumably biallelic KO), that were only BFP expressors but with two different populations of higher and lower levels of expression (presumably with one or two alleles with KI) and that co-expressed GFP and BFP (presumably one allele each) **(Fig. 1B)**. Using this phenotyping strategy, the edition rate in the GFP+ BFP- subpopulation is underestimated because GFP/GFP and GFP/KO (frameshift mutation) cells could neither be discriminated from each other nor the generation of indels that do not eliminate the fluorescente activity of the GFP (in frame mutation).

**Figure 1:**
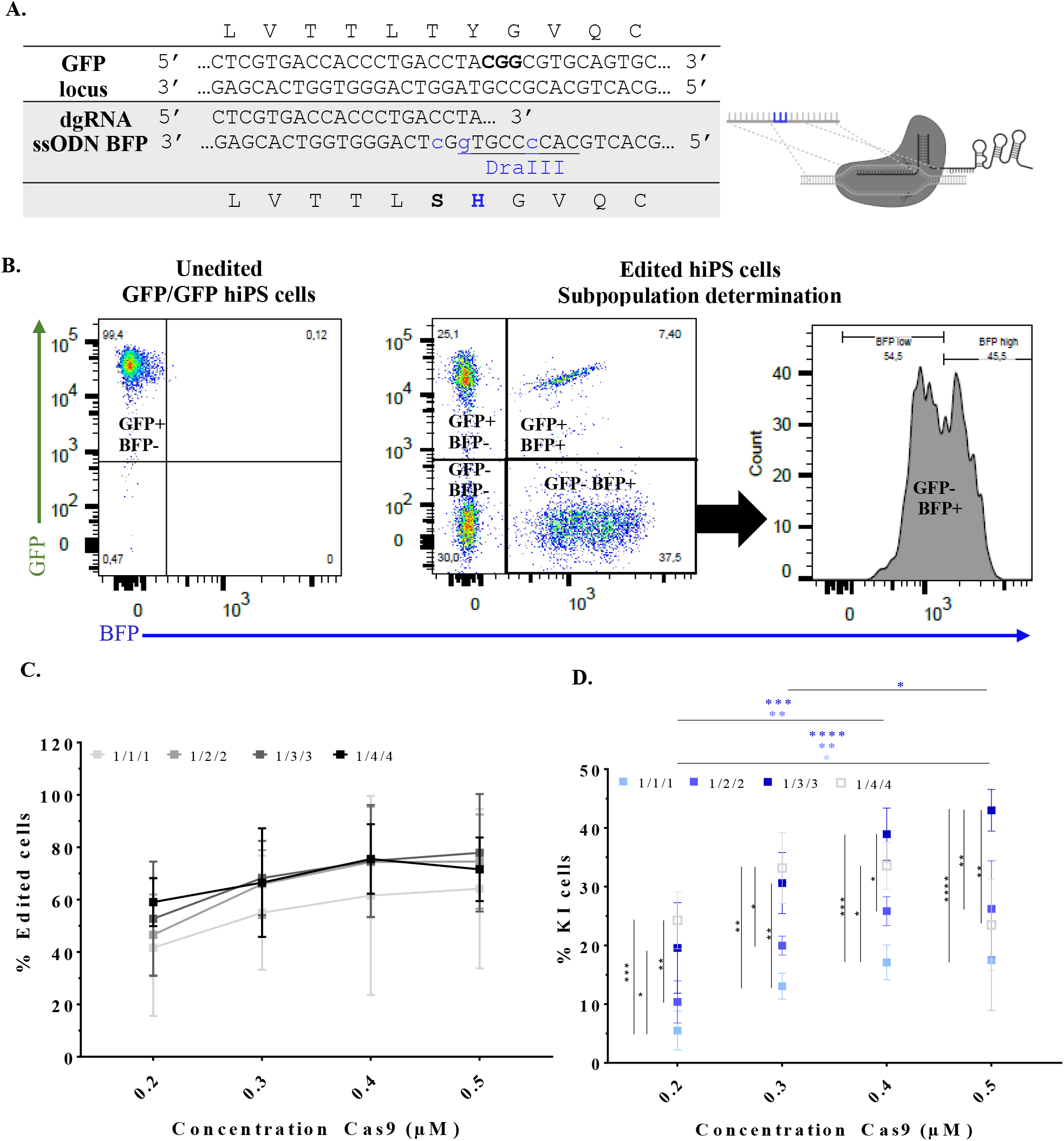
Molecular optimization of GFP to BFP conversion on hiPS cells. **A**. Schematic representation of GFP to BFP conversion strategy using CRISPR/Cas9 and a BFP ssODN. PAM is written in bold, KI in blue and DraIII restriction site is underlined. **B**. Discrimination of outcomes after CRISPR/Cas9 edition with BFP ssODN on GFP hiPS cells using flow cytometry phenotyping. GFP/KO cells cannot be discriminated from GFP/GFP cells using this strategy. **C**. Effect of increased concentrations of Cas9 protein on edition rate (include KI) using Cas9/dgRNA/ssODN 1/1/1, 1/2/2, 1/3/3 and 1/4/4 ratios. **D**. Effect of increased concentrations of Cas9 protein on KI rate using the same ratios. Asterisks highlight the statistical differences between conditions using two ways ANOVA (*P≤ 0.05, **P≤0.01, ***P≤0.001 and ****P ≤ 0.0001). Figure created with BioRender.com. GFP, Green Fluorescent Protein; dgRNA, dual guide RNA; ssODN, single-stranded oligonucleotide; BFP, Blue Fluorescent Protein; hiPS, human induced pluripotent stem cells.

Skarnes et al. electroporated 800,000 hiPS cells in 100μL with 20μg of Cas9 (1.2μM), 16μg (5μM) of gRNA and 200pmol (2μM), Cas9/sgRNA/ssODN 1/5/2 molar ratio using Amaxa 4D nucleofector (Lonza) pulse code CA-137 (Skarnes et al. 2019). For convenience, we decided to also use this electoporator but with 200,000 cells in 20μL. For a starting point, we challenged our GFP expressing hiPS with 0.2μM or 1μM of Cas9 protein with a Cas9/dgRNA/ssODN ratio of 1/1/1 and 0.5μM or with a 1/5/5 ratio. Cells electroporated with 1μM of Cas9 1/1/1 ratio (15% of confluence) display about 2 times more toxicity than those with 0.2μM at the same ratio (35% of confluence), preventing us from ratio increase with this condition **(Fig. S1A)**. Moreover, we observed even higher toxicity with cells electroporated with 0.5μM 1/5/5 ratio (5% of confluence) indicating that this ratio is out of the limit that these cells can bear **(Fig. S1A)**. That is why we decided not to repeat these conditions and to limit Cas9 concentrations range to 0.2, 0.3, 0.4 and 0.5μM and Cas9/dgRNA/ssODN ratio to 1/1/1, 1/2/2, 1/3/3 and 1/4/4. Edition rates (including KO and KI) with 1/1/1 ratio tend to improve with increased doses of Cas9 protein from 0.2 μM to 0.5 μM (41.7% and 64.2% respectively) but with no significant differences **(Fig. 1C)**. The same trend is observed for all ratios except for 1/4/4 with 0.5μM of Cas9 protein. For this condition and 0.4μM of Cas9 1/4/4 ratio, we observed less than 5% of confluence two days after electroporation while having more than 25% of confluence for 1/1/1 ratio with the same Cas9 concentrations (**Fig. S1B**), which indicates that the higher limits of doses of dgRNA and ssODN we can use have been crossed. The edition rates are similar between ratios for each concentration of Cas9 tested.

For Cas9/dgRNA/ssODN 1/1/1 ratio, KI rates were increased significantly with higher doses of Cas9 from 0.2 μM to 0.5 μM (5.5% and 17.5% respectively, p=0.0471) **(Fig. 1D)**. This difference between 0.2 and 0.5μM of Cas9 on KI is also observed with 1/2/2 (10.4% and 26.2% respectively, p=0.0012) and 1/3/3 (19.6% and 43.0% respectively, p<0.0001) ratios. KI rate with 0.4μM of Cas9 is significantly higher than 0.2μM for 1/2/2 ratio (25.8% and 10.4% respectively, p=0.0035) and 1/3/3 (39.0% and 19.6% respectively, p=0.0006). These data confirm that 0.2μM of Cas9 is too low for efficient KI and that 0.4μM could be enough. In the same way, increasing Cas9/dgRNA/ssODN ratios significantly improved KI rates. For 0.4μM of Cas9, 1/1/1 KI rate (17.1%) is significantly lower than 1/3/3 and 1/4/4 (39.0% p=0.0005 and 33.6% p=0.0103 respectively). Nevertheless, for 0.5μM of Cas9, 1/4/4 KI rate is significantly lower than 1/3/3 (23.5% and 43.0% respectively, p=0.002) and could be explained by toxicity (**Fig S1B**), as discussed for edition. Thus, 0.4μM Cas9 protein seems a good compromise between efficacy and toxicity. The toxicity observed for 0.5μM of Cas9 1/4/4 ratio may be due to high amount of dgRNA not loaded to Cas9 protein. The fact that one Cas9 can load only one dgRNA suggest that, if RNP complex formation is efficient, RNP (Cas9/dgRNA) ratio may be reduced to use high concentrations of ssODN (2μM tested here) with limited toxicity.

### Assessment of RNP complex formation efficiency *in vitro* by NanoDSF

RNP complex formation is characterized by a conformational rearrangement which affect Tm and could be detected by NanoDSF. To define the efficacy of RNP complex formation and the best ratio of dgRNA per protein of Cas9, we obtained the denaturation curve induced by temperature increase. Using the first derivative, we determined the melting temperature (Tm) of Cas9 alone (Tm_Cas9_) or with increased concentrations of dgRNA (Tm_RNP_) matching RNP ratios electroporated into ratmbryos and hiPS cells (**Fig. 1A** and **1B**, respectively). In all conditions, addition of dgRNA rEphx2, used for the genome editing of rat embryos **(Fig. 2A)**, or dgRNA GFP, used in hiPS genome editing strategy **(Fig. 2B)**, induced a drastic Tm increase compared to Cas9 alone and reflects RNP complex formation. For molar RNP (Cas9/dgRNA) ratios rEphx2 in favor of Cas9 (RNP 5/1 and 2/1 ratios) we observed a denaturation curve with two transitions represented as Tm_Cas9_ and Tm_RNP_ **(Fig. 2A)**. Compared to Cas9 alone (Tm_Cas9_ Cas9=43.1°C), Tm_Cas9_ for those two RNP ratios were not signicantly different (Tm_Cas9_ RNP 5/1=42.9°C; Tm_Cas9_ RNP 2/1=42.9°C, p=0.4809 and 0.9353, respectively) while Tm_RNP_ were (Tm_RNP_ RNP 5/1=47.8°C; Tm_RNP_ RNP 2/1=48.7°C, p=0.029 for both). These results indicate that uncomplexed Cas9 remains in these conditions. With an equal ratio (RNP 1/1) or ratios in favor of dgRNA (RNP 1/2, 1/3 and 1/5), we observed only one transition, (Tm_RNP_ RNP 1/1= 49.8°C; Tm_RNP_ RNP 1/2 =49.8°C; Tm_RNP_ RNP 1/3=49.6°C; Tm_RNP_ RNP 1/5=49.7°C) which were not significantly different from each others. Moreover, TmRNP for RNP 1/1 ratio is sigificantly different from RNP 2/1 ratio (Tm_RNP_ RNP 2/1 = 48.7°C; Tm_RNP_ RNP 1/1=49.8°C, p=0.0206) and Cas9 alone (Tm_Cas9_ Cas9=43.1°C; Tm_RNP_ RNP 1/1=49.8°C, P<0.0001). This indicate that all Cas9 have formed an RNP complex with RNP 1/1 and ratios in favor of dgRNA. TmRNP RNP 1/1 with dgRNA hiPS GFP is also significantly different from Cas9 alone (Tm_Cas9_ Cas9=43.1°C; Tm_RNP_ RNP 1/1=49.5°C, P=0.0029) and ratios in favor of gRNA (RNP 1/2, 1/3 and 1/5) are not sificantlty different from each others either. Thus, RNP complex formation is efficient in the conditions used *in vitro* and RNP 1/1 ratio seems optimal for *in vivo* application.

**Figure 2:**
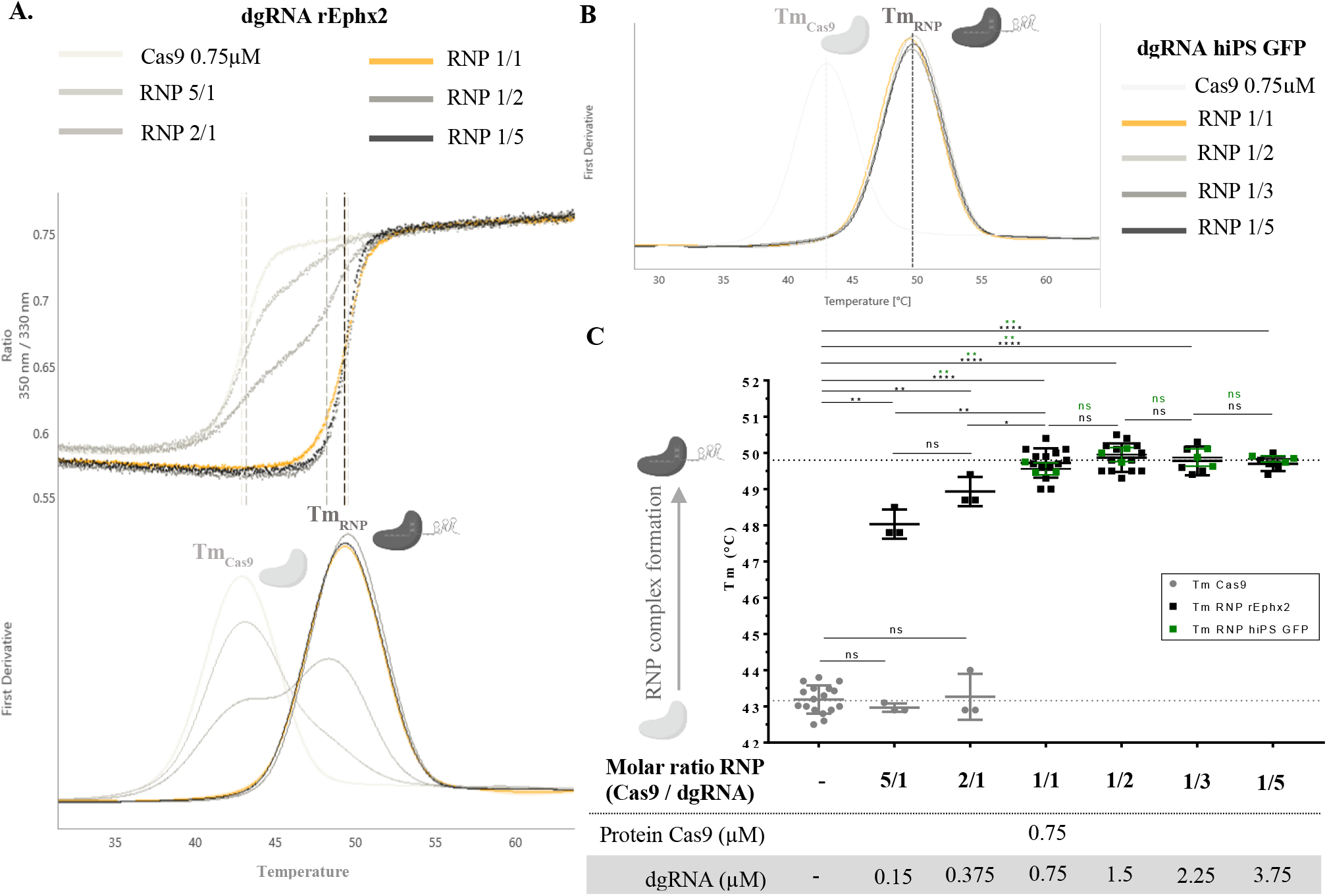
Assessment of RNP complex formation efficiency *in vitro* by NanoDSF. **A**. NanoDSF curves and derivatives used to determine Tm of Cas9 (Tm_Cas9_) and RNP complex (Tm_RNP_) for each condition tested with dgRNA rEphx2. **B**. NanoDSF derivatives used to determine Tm_Cas9_ and Tm_RNP_ with dgRNA hiPS GFP. **C**. Comparison of RNP complex formation efficiency by Tm_Cas9_ and Tm_RNP_ analysis with increased concentrations of both dgRNA. Asterisks highlight the statistical differences between conditions using nonparametric Mann-Whitney tests (nsP>0.05, *P≤0.05, **p ≤ 0.01, ***p ≤ 0.001 and ****p ≤ 0.0001). Figure created with BioRender.com. dgRNA, dual guide RNA; RNP, ribonucleoproteic complex; Tm, melting temperature; hiPS GFP, human induced pluripotent stem cells expressing the Green Fluorescent Protein.

### Highly efficient hiPS GFP to BFP conversion and characterization

By phenotyping, we compared rates of KI and genome edition (KI and KO) but also the proportion of each subpopulation generated by electroporation of hiPS cells with 0.4μM of Cas9, RNP 1/1 or 1/2 ratios and 2μM of ssODN **(Fig. 3A)**. Increase of ssODN concentrations to 3 or 4μM lead to higher toxicity **(Fig. S2)**. We achieved similarly high KI (RNP 1/1=55.4%; RNP 1/2=54.9%) and edition (RNP 1/1=83%; RNP 1/2=84.4%) efficiencies between RNP 1/1 and 1/2 ratios. Moreover, RNP 1/1 and 1/2 ratios generated not significantly different proportions of GFP+ BFP- (RNP 1/1=18.0%; RNP 1/2=15.0%), GFP+ BFP+ (RNP 1/1=4.9%; RNP 1/2=2.5%), GFP- BFP high (RNP 1/1=23.3%; RNP 1/2=24.6%), GFP- BFP low (RNP 1/1=27.2%; RNP 1/2=27.8%) and GFP- BFP- (RNP 1/1=27.6%; RNP 1/2=29.5%).

**Figure 3:**
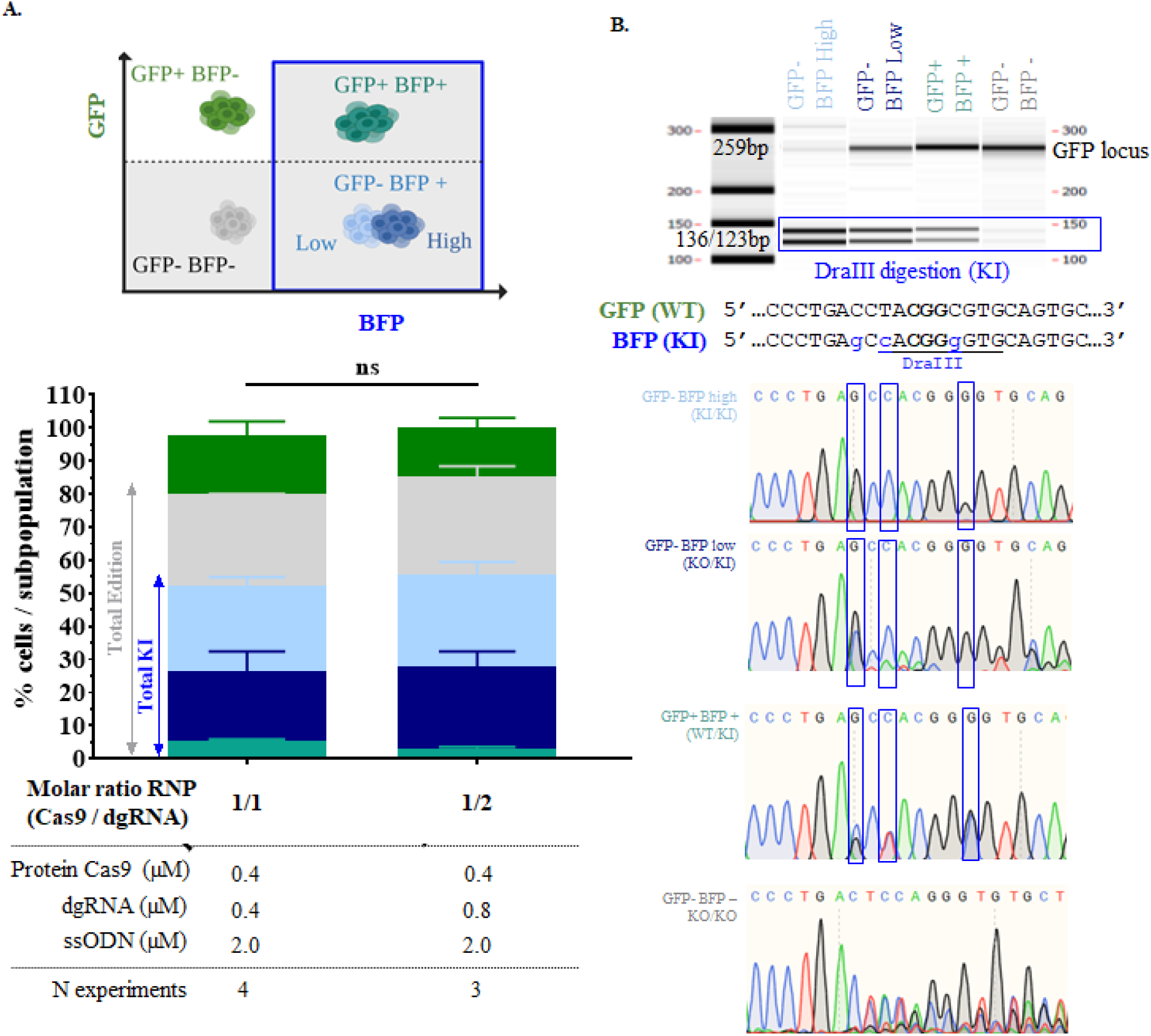
Highly efficient hiPS GFP to BFP conversion and characterization. **A**. Comparison of KI, edition rate and proportions of each subpopulation (defined by phenotyping) between RNP (Cas9/dgRNA) 1/1 and 1/2 ratios with 2μM of ssODN BFP. Two ways ANOVA test was used to verify the significance of the difference between the two conditions tested (nsP>0.05). **B**. Subpopulation genotyping by DraIII digestion and Sanger sequencing of previously sorted subsets. DraIII was performed on GFP amplicon (259bp) which amplified WT, indels and KI alleles. DraIII digestion of KI allele generated two bands (136 and 123bp) on capillary electrophoresis. Sequencing was performed on GFP amplicon and compared to GFP and BFP sequence. BFP mutation are noted in blue, DraIII restriction site is underlined and dgRNA hiPS GFP is written in bold. Figure created with BioRender.com. GFP, Green Fluorescent Protein; BFP, Blue Fluorescent Protein; dgRNA, dual guide RNA; ssODN, single-stranded oligonucleotide; WT, Wild-type; KI, knockin; KO, knockout.

These results indicate that 0.4μM of Cas9, with RNP 1/1 ratio and 2μM of ssODN, is sufficient and optimal to reach high efficiency of KI generation in hiPS cells. Using cell sorting **(Fig. S3)** we genotyped the subpopulations generated by electroporation of this optimal condition by genotyping. DraIII digestion **(Fig. 3B)**, which reflects BFP conversion by ssODN mediated KI, was consistent with Sanger sequencing **(Fig. 3B)**. GFP- BFPhigh cells profil analysis showed full DraIII digestion and only BFP sequence, demonstrating their expected KI/KI genotype. GFP- BFPlow cells display partial (67.6%) DraIII digestion with BFP sequence coupled with various indels profiles on sequencing suggesting a KO/KI genotype. GFP+ BFP+ analysis showed also partial (32%) DraIII digestion and both GFP and BFP sequences, confirming the expected WT/KI genotype. Sample with GFP- BFP- cells was digested in an extremely low proportion by DraIII (digested bands under the quantifiable range of the system). This sequence indicated diverse indels profiles and no BFP sequence, confirming the expected KO/KO genotype. The coherence between phenotypes and genotypes of CRISPR/Cas9 outcomes on the GFP to BFP conversion model validate the power of this phenotyping technic.

### Highly efficient point mutations on rat embryos

To define optimal conditions of KI in one-cell rat embryos, an XbaI restriction site was generated by point mutation on the *rEphx2* locus using CRISPR/Cas9 and ssODN electroporation **(Fig. 4A)**. KI and edition rats were assessed by an heteroduplex mobility assay on microfluidic capillary electrophoresis (HMA-CE) (Chenouard et al. 2016), Xba I digestion, sequencing and quantitative polymerase chain reaction (qPCR). Increasing Cas9 protein concentration from 0.1μM to 0.2μM drastically improved both edition (73.1% and 96.8%, respectively) and KI (30.8% and 51.6%, respectively) rates with RNP 1/1 ratio and 2μM of ssODN **(Fig. 4B and Table S1)**. On the other hand, increasing Cas9 protein concentration to 0.4μM reduced KI efficiency (34.2%). RNP 1/2 ratio with 0.2μM of Cas9 also reduced KI rate (RNP1/1=51.6%; RNP1/2=37.5). Toxicity for all conditions was comparable to the usual results obtaines following microinjection or unmanipulated rat embryos, with more than 25% of birth (table S1). Thus, electroporation of 0.2μM of Cas9 protein with ratio RNP1/1 and ssODN 2μM was the optimal condition to achieve high KI efficiency on rat embryos.

**Figure 4:**
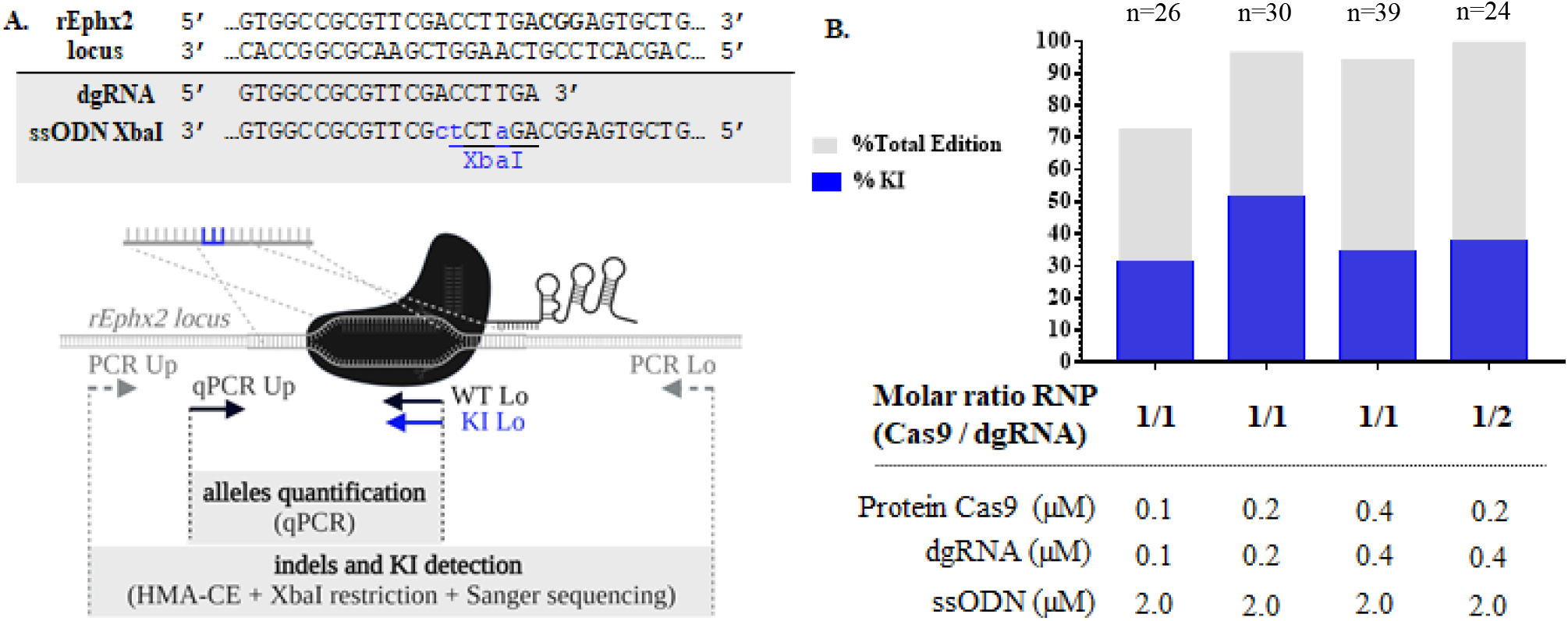
Highly efficient point mutations on rat embryos. **A**. Schematic representation of rEphx2 point mutation generation by CRISPR/Cas9 and genotyping strategies. PAM is written in bold, KI in blue and DraIII restriction site is underlined. **B**. Highly efficient KI generation on rEphx2 locus using low doses of CRISPR Cas9. Figure created with BioRender.com. dgRNA, dual guide RNA; ssODN, single-stranded oligonucleotide; WT, Wild-type; KI, knockin; qPCR, quantitative polymerase chain reaction; HMA-CE, heteroduplex mobility assay by capillary electrophoresis.

## III. Discussion

Methodologies for application of CRISPR/Cas9 to cells or embryos are very diverse between laboratories in particular concentrations of Cas9 protein, gRNA and donor DNA. Those concentrations are usually expressed in mass and not in mole, which makes it harder to compare different works and extrapolate the molecular rational for each component. In our study, we suggest to apply molecular rational on CRISPR/Cas9 for genome editing setup. To do so, we developped a technic to verify RNP complex formation efficiency. Indeed, if one Cas9 can only load one gRNA, optimal RNP (Cas9/dgRNA) ratio should then be close to 1/1 for efficient conditions of preparation of RNP complex. Nevertheless, the use of single or dgRNA as well as the donor DNA matching the guide RNA designed for KI may modify this interaction with Cas9 and need to be tested in future experiments for each specific gene editing experiment. DSF has been previously used by two teams to qualitatively charaterize Cas9 thermostability in the presence of truncated or unfolded gRNA (Tillotson 2017) or in the presence of different DNA target binding conditions (Jiang et al. 2016). We have also tested this technique (data not shown) but the requirement of a dye make it poorly reproducible. NanoDSF is a label and immobilization-free technique that allows very high sensibility and reproducibility. With this method we can highlight the modification of thermal stability of the Cas9 protein induced by its complexation with dgRNA. This assay emphasis the importance of optimal RNP complex formation for efficient edition and KI, eventhough it is rarely studied in the literature. Futher experiments need to be performed to explore other guide RNAs and different formats.

This rational molar optimization of Cas9 and dgRNA prevented excess of free dgRNA to limit toxicity and allowed the use of higher ssODN concentrations *in vivo*. It facilitated KI model generation both on iPS cells and rat embryos by defining optimal conditions as 0.4μM and 0.2μM of Cas9 protein respectively, with RNP 1/1 ratio and 2μM of ssODN. Using these conditions, we reached more than 50% of KI for both models while greatly reducing our costs for reagents. Our study demonstrate that it is not necessary to use neither high amount of Cas9 nor high RNP (Cas9/gRNA) ratio to achieve high KI rate compared to previously published conditions in hiPS cells (Skarnes et al. 2019; Xu et al. 2019) and rats (Kaneko and Nakagawa 2020; Remy et al. 2017) by us and others. This optimization will accelerate the development of KI models using CRISPR/Cas9. In particular for the rat, it will reduce the number of embryos necessary to develop a model and more easily tends toward 3Rs. Moreover, this study may pave the way for more efficient KI with long DNA donor by increasing the chance of DNA donor occurence at Cas9 cut site with rational molar approach.

## IV. Material and Methods

### CRISPR/Cas9 components and RNP formation

Cas9 protein, crRNA and tracrRNA were purchased from Integrated DNA Technologies (Alt-R^®^ CRISPR Cas9 system) as well as ultramer ssODN modified on both extremities with phosphorothioate bonds. Sequences are specified in table S2.

Dual gRNA is formed by equimolar incubation of crRNA and tracrRNA in Nuclease Free Duplex Buffer (30 mM HEPES, pH 7.5; 100 mM potassium acetate) from Integrated DNA Technologies. It is denatured 5minutes at 95°C and incubated 5min at 25°C to ensure correct folding of dgRNA. RNP complex is formed by incubation of Cas9 protein and dgRNA 10 minutes at 25°C in HEPES 20mM, KCl 150mM, pH7,5 buffer. Concentrations used are specified for each experiment. ssODN is added at concentrations specified for each experiment after RNP formation.

### NanoDSF

10 μL of RNP complex prepared with 0.75 μM of Cas9 were loaded into nanoDSF grade high sensitivity capillaries (NanoTemper Technologies) and installed on capillary array on the Prometheus NT.48 NanoDSF instrument (NanoTemper Technologies). The temperature was increased from 20 °C to 95 °C with a linear thermal ramp (at a rate of 1 °C/min). Changes in fluorescence of tryptophan at 330 and 350 nm were recorded at a rate of 10 datapoints per minute. The 350/330 nm ratio of the two wavelengths were plotted against temperature and the first derivative analysis allowed the determination of melting temperatures (Tm) using the PR.Therm Control Software.

### hiPS cells

#### Culture of hiPSC and generation of GFP iPSC lines

iPSC were cultured in feeder-free culture conditions: stem cell-qualified Matrigel-coated plates (0.1 mg/mL; BD Bioscience) with mTeSR (Stem Cell Technologies) and passaged using the Passaging Solution XF (StemMACSTM, Miltenyi Biotec). When using single cells suspension, cells were detached with TrypLE^™^ Express Enzyme (Thermofisher Scientific _ 12605010) and culture medium was supplemented with 10μM Rock inhibitor Y27632 (Sigma_ 104M4646V).

For the GFP expressing lines, 10^6^ human induced pluripotent stem cells “LON71-019” (Gaignerie et al. 2018) were transfected with 3μg TALEN R + 3μg TALEN L + 4.8μg AAV-CAGGS-EGFP (addgene #22212) using Amaxa^™^ Nucleofector^™^ II (P3 kit, B-16 program) to obtain KI in the AAV locus. Clones were manually selected and first maintained on matrigel-coated plates with Stemflex, then switched to mTeSR1 medium. Genomic PCR of the AAV locus allowed to select cells that were bi-allelically edited. Genomic integrity of edited lines was performed with Integragen SNP.

#### GFP to BFP edition

Gentle TryplE (Gibco) enzymatic digestion was performed to obtain single cells suspension. 200k cells, resuspended in P3 Primary Cell NucleofectorTM Solution (Lonza), were mixed with RNP complex and the ssODN that contains the sequence that will edit GFP to BFP sequences as well as a new DraIII restriction site (Glaser et al. 2016) followed by electroporation in 16-well Nucleocuvette^®^ Strips (Lonza) using CA-137 program. Seven days after transfection, cells were dissociated using TryplE. Dead cells were stained with Fixable Viability Stain 440UV (BD biosciences). Cells were analyzed using BD FACSCelesta^™^ (GFP: laser: 488nm, filter: 530/30; BFP: 405nm laser: 450/40, Viability: laser Ultraviolet, filter: 379/28) and sorted with BD FACSAria^™^ (GFP: laser: 488nm, filter: 530/30; BFP: 405nm laser: 450/40).

#### Animals

Sprague-Dawley (SD) rats were the only strain used and were sourced from Janvier Labs (Le Genest-Saint-Isle, France). All the animal care and procedures performed in this study were approved by the Animal Experimentation Ethics Committee of the Pays de la Loire region, France, in accordance with the guidelines from the French National Research Council for the Care and Use of Laboratory Animals (Permit Numbers: CEEA-PdL-2020-26567).

#### Electroporation of rat intact zygotes and embryo transfer

The electroporation procedure and embryo transfer were performed as previously described (Remy et al. 2017). Briefly, fertilized 1-cell stage embryos were collected from super-ovulated prepubescent females (30IU PMSG at day-2 before breeding and 20IU HCG the day of breeding; Intervet, France) and kept in M16 medium at 37°C under 5% CO2 until electroporation.

Electroporation with dgRNA/Cas9 RNP complex was performed using the NEPA21 system (NEPA GENE Co. Ltd, Sonidel, Ireland) at 300V 0,5ms to limit toxicity. Surviving embryos were implanted the same day in the oviduct of pseudo-pregnant females (0.5dpc) and allowed to develop until embryonic day 14 (E14).

#### Genotyping

Briefly, hiPS cells and E14 embryos were digested overnight at 56°C in 1mL of tissue digestion buffer (Tris-HCl 0.1 mol/L pH 8.3, EDTA 5 mmol/L, SDS 0.2%, NaCl 0.2 mol/L, PK 100 μg/mL). The CRISPRs nuclease targeted regions were PCR amplified from diluted lysed samples (1/20) with a high-fidelity polymerase (Herculase II fusion polymerase). To detect gene edition, locus specific primers *GFP* and *rEphx2* (Table S2) were used and mutations were analyzed using a heteroduplex mobility microfluidic capillary electrophoresis assay as previously described in detail (Chenouard et al. 2016) followed by PCR amplicon direct digestion by XbaI and sequencing. qPCR was done with primers described in table S2 to quantify both WT and KI alleles.

#### Statistical test

Graphs and statistical analyses were generated using GraphPad Prism v5.03. Two ways ANOVA tests were performed to compare KI rate, edition rate and subpopulation proportion on hiPS GFP locus. NanoDSF Tm comparison were performed using nonparametric Mann-Whitney tests.

## Acknowledgments

We thank all our colleagues from genOway, CRTI and the TRIP plateform for their support. In particular we thank Claire Usal and Séverine Ménoret for their animal expertise, Laure-Hélène Ouisse, Ghenima Ahmil, Séverine Bézie and Jenny Greig for their flow cytometry and cell sorting expertise, Aude Guiffes for her work on rat genotyping and Sarah Duponchel for her wise advices. NanoDSF studies was supported by funding from the Biogenouest network and from the “Pays de la Loire” region.

## Author contributions

VC participated in all experiments design and analysis. IL and LD designed hiPS cells experiments and IL performed experiments and analysis. SR and IL phenotyped hiPS cells by flow cytometry. VC performed RNP complex formation for rat embryo electroporation and NanoDSF studies. SR handled rat embryos and electroporation of RNP complex. LT set up and performed hiPS cells and rat embryos genotyping. AF and KB performed NanoDSF experiments and analysis. YC and IA secured funding. IA defined experiments and interpreted data. VC wrote this manuscript with all authors intellectual contribution.

