## Supplementary material for "Highly Efficient Knockin in Human iPS Cells and Rat Embryos by CRISPR/Cas9 Molecular Optimization": Fig. S1

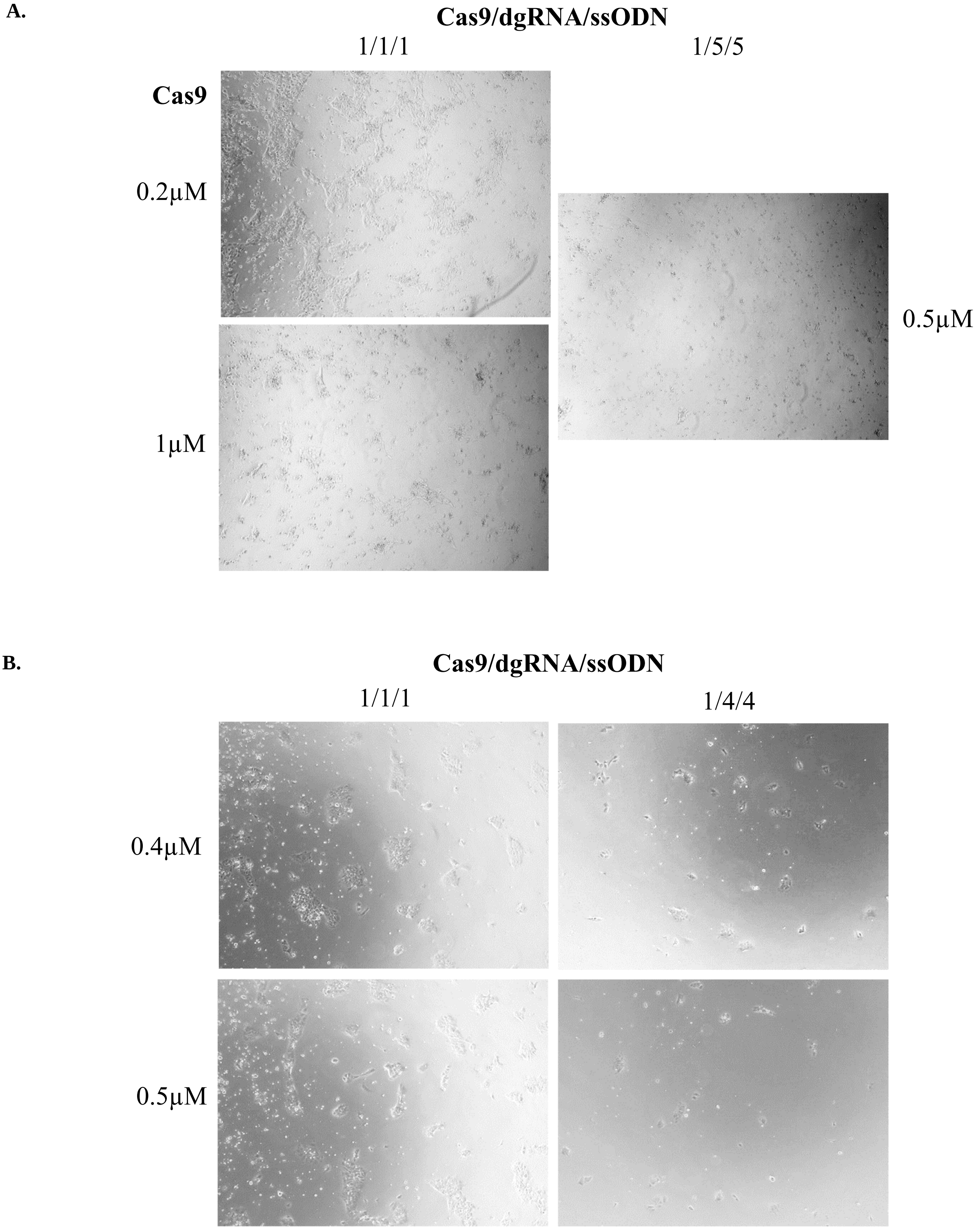


**Figure S1: Observation of hiPS cells culture and formation of colonies with Zeiss Axio vert.A1 microscope (5X) 2 days after electroporation of Cas9 protein with ratio Cas9/dgRNA/ssODN. A.** Cells edited with 0.2µM of Cas9 protein ratio Cas9/dgRNA/ssODN 1/1/1 formed nice colonies indicating that they have well recovered from electroporation 2 days later (35% of confluency). In comparison, cells electroporated with 0.5µM ratio 1/5/5 and 1µM of Cas9 with 1/1/1 display much more toxicity (about 5 and 15% of confluency, respectively) and form less and smaller colonies. **B.** Cells electroporated with ratio 1/4/4 and 0.4 and 0.5µM of Cas9 showed 5 times more toxicity (5% and 3% confluency, respectively) than the same concentration of Cas9 with ratio 1/1/1 (25% of confluency for both conditions). Ratio 1/1/1 allow formation of numerous colonies compared to higher ratios. dgRNA, dual guide RNA; ssODN, single-stranded oligonucleotide.
