## Supplementary material for "Highly Efficient Knockin in Human iPS Cells and Rat Embryos by CRISPR/Cas9 Molecular Optimization": Fig. S2

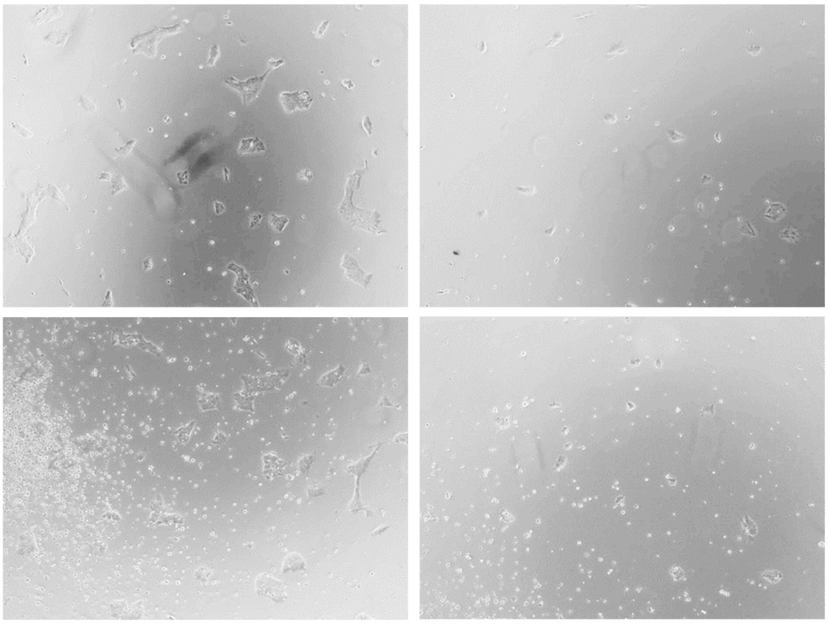


**Figure S2: Microscopic observation with Zeiss Axio vert.A1 microscope (5X) of hiPS cells 2 days post-electroporation with Cas9 0.4µM, RNP ratio Cas9/dgRNA 1/1 and 2, 3 or 4µM of ssODN.** Electroporation of 3µM and 4µM of ssODN with Cas9 0.4µM and RNP ratio 1/1 shows low confluences (10% and 5% respectively compared to their 2µM control (25% and 20%, respectively). Moreover, with 2µM of ssODN, hiPS cells form numerous colonies compared to higher concentrations. dgRNA, dual guide RNA; ssODN, single-stranded oligonucleotide.
