## Supplementary material for "Highly Efficient Knockin in Human iPS Cells and Rat Embryos by CRISPR/Cas9 Molecular Optimization": Fig. S3

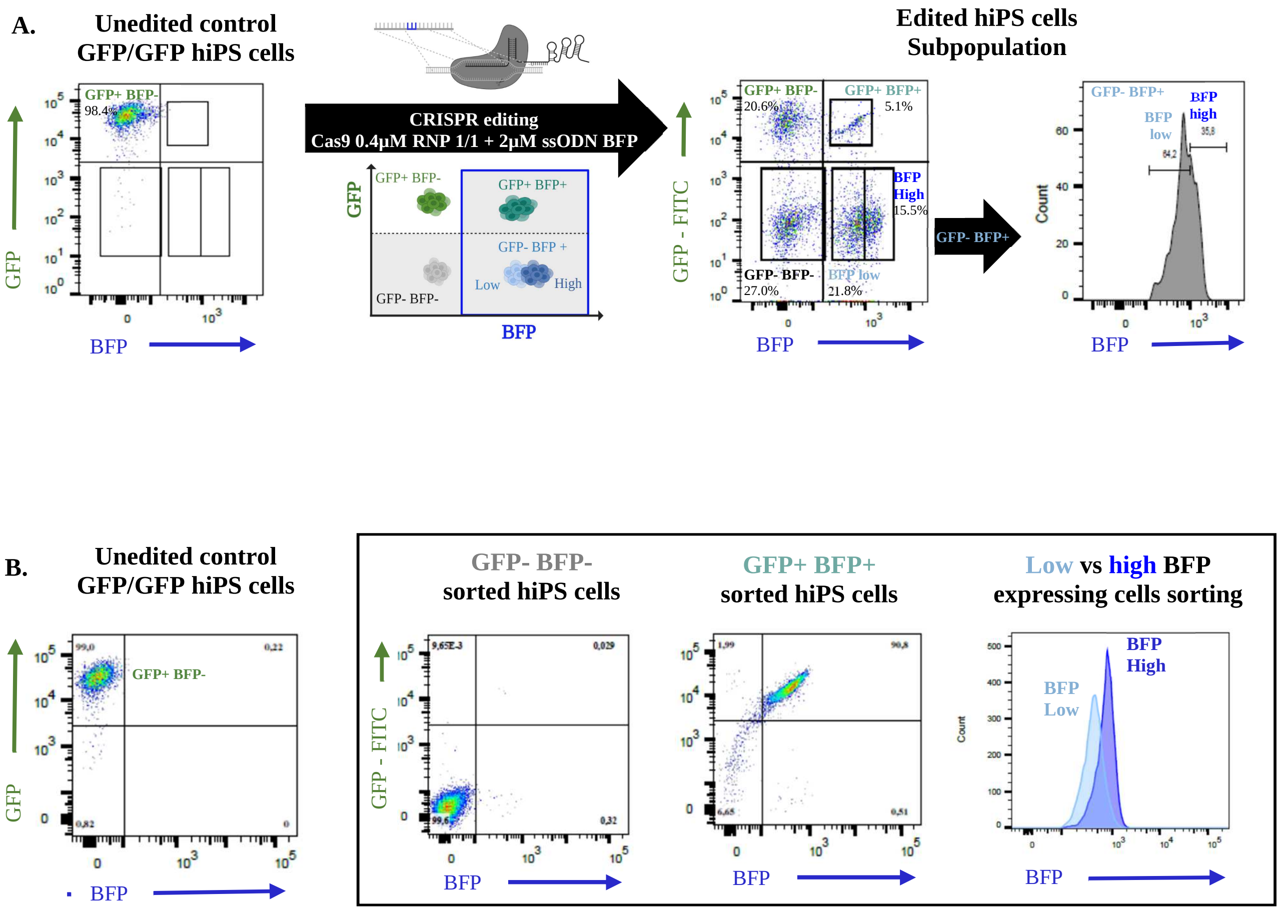


**Figure S3: Cell sorting strategies to discriminate subpopulations obtained by CRISPR/Cas9 edition with 0.4µM of Cas9 RNP (Cas9/dgRNA) 1/1 ratio and 2µM of ssODN BFP.** Phenotyping of hiPS cells GFP expressing cells edited with 0.4µM of Cas9 RNP (Cas9/dgRNA) ratio 1/1 and 2µM of ssODN before **A.** and after **B.** cell sorting of GFP- BFP-, GFP+ BFP+, GFP- BFP low and GFP- BFP high subpopulations. hiPS, human induced pluripotent stem cells; GFP, Green Fluorescent Protein; BFP, Blue Fluorescent Protein; ssODN, single-stranded oligonucleotide.
