## Supplementary material for "Highly Efficient Knockin in Human iPS Cells and Rat Embryos by CRISPR/Cas9 Molecular Optimization": Table S1

**Table S1. CRISPR/Cas9 mediated edition and KI generation on rat embryos with dgRNA rEphx2 and ssODN XbaI.**

| Cas9 (µM) | Ratio RNP (Cas9/dgRNA) | ssODN (µM) | No. of days of electroporation | No. of foster females used | No. of analyzed /No. of transferred embryos (%) | No. edited embryos, include KI (%) | No. KI embryos (%) |
|---|---|---|---|---|---|---|---|
| 0.1 | 1/1 | 2 | 4 | 4 | 26 / 104 (25.0%) | 19 (73.1%) | 8 (30.8%) |
| 0.2 | 1/1 | 2 | 3 | 4 | 30 / 108 (27.8%) | 29 (96.8%) | 16 (51.6%) |
| 0.4 | 1/1 | 2 | 3 | 4 | 39 / 115 (33.9%) | 37 (94.9%) | 13 (34.2%) |
| 0.2 | 1/2 | 2 | 3 | 3 | 24 / 88 (27.3%) | 24 (100%) | 9 (37.5%) |

ssODN, single-stranded oligonucleotide; KI, knockin.
