## Supplementary material for "Highly Efficient Knockin in Human iPS Cells and Rat Embryos by CRISPR/Cas9 Molecular Optimization": Table S2

**Table S2. dgRNA, ssODN and genotyping primers sequences.**

| Reactifs | Name | Sequence (5'-3') | Ref. |
|---|---|---|---|
| **gRNA** | rEphx2 2.3 | GTGGCCGCGTTCGACCTTGA | (Remy et al. 2017) |
| | EGFP 2 | CTCGTGACCACCCTGACCTA | (Glaser et al. 2016) |
| **ssODN** | rEphx2 KI XbaI | *G*C*T*GCAGCCCGGGCCATCATGGCGCTGCGTGTGGCCGCGTTCGctCTaGACGGAGTGCTGGCCCTCCCCTCTATAGCCGGGGTTCTGCGCCACACC*G*A*G*G | (Remy et al. 2017) |
| | ssODN BFP2 antisense | *G*G*C*ATGGCGGACTTGAAGAAGTCGTGCTGCTTCATGTGGTCGGGGTAGCGGCTGAAGCACTGCACCCCGTGGCTCAGGGTGGTCACGAGGGTGGGCCAGGGCACGGGCAGCTTGCCGGTGGTGCAGATGAACTTC*A*G*G*GT | (Glaser et al. 2016) |
| **HMA-CE primers** | rEphx2 Up | GGCAGGGTTTCTAGTTCTTGG | (Remy et al. 2017) |
| | rEphx2 Lo | TCTTGTAAACTGAGGCGGGTA | (Remy et al. 2017) |
| | rEphx2-LongUp | ACCTCATCATTCCTTCCTCAGTT | |
| | rEphx2-Lo2 | GTGACTGGAGGCGATGTTGT | |
| | rEphx2-Lo4 | CACGGCAACACCAGAGCTTA | |
| | GFP-Up1 | CGTAAACGGCCACAAGTTCA | |
| | GFP-Lo4 | CTTGTAGTTGCCGTCGTCCT | |
| **qPCR primers** | rEphx2-188For | GGCGCTGCTCTAGTCTTAGGTTT | |
| | rEphx2-WT16Rev | AGCACTCCGTCAAGGT | |
| | rEphx2-KI16Rev | AGCACTCCGTCTAGAG | |
| | rAnks3-205For | CCCCAGCCTCCCACTTGTC | |
| | rAnks3-205Rev | AGGATGACTGAAATTGGTGGAGTTGC | |

*indicates bases with phosphorothioate modification. ssODN, single-stranded oligonucleotide; KI, knockin, qPCR, HMA-CE, heteroduplex mobility assay by capillary electrophoresis; quantitative polymerase chain reaction; GFP, Green Fluorescent Protein; BFP, Blue Fluorescent Protein; hiPS, human induced pluripotent stem cells.
